# Co-estimating Reticulate Phylogenies and Gene Trees from Multi-locus Sequence Data

**DOI:** 10.1101/095539

**Authors:** Dingqiao Wen, Luay Nakhleh

**Affiliations:** Department of Computer Science, Rice University, 6100 Main Street, Houston, TX 77005, USA; Department of BioSciences, Rice University, 6100 Main Street, Houston, TX 77005, USA

**Author notes:** Correspondence to be sent to: Department of Computer Science, Rice University, 6100 Main Street, Houston, TX 77005, USA;.

## Abstract

The multispecies network coalescent (MSNC) is a stochastic process that captures how gene trees grow within the branches of a phylogenetic network. Coupling the MSNC with a stochastic mutational process that operates along the branches of the gene trees gives rise to a generative model of how multiple loci from within and across species evolve in the presence of both incomplete lineage sorting (ILS) and reticulation (e.g., hybridization). We report on a Bayesian method for sampling the parameters of this generative model, including the species phylogeny, gene trees, divergence times, and population sizes, from DNA sequences of multiple independent loci. We demonstrate the utility of our method by analyzing simulated data and reanalyzing three biological data sets. Our results demonstrate the significance of not only co-estimating species phylogenies and gene trees, but also accounting for reticulation and ILS simultaneously. In particular, we show that when gene flow occurs, our method accurately estimates the evolutionary histories, coalescence times, and divergence times. Tree inference methods, on the other hand, underestimate divergence times and overestimate coalescence times when the evolutionary history is reticulate. While the MSNC corresponds to an abstract model of “intermixture,” we study the performance of the model and method on simulated data generated under a gene flow model. We show that the method accurately infers the most recent time at which gene flow occurs. Finally, we demonstrate the application of the new method to a 106-locus yeast data set. [Multispecies network coalescent; reticulation; incomplete lineage sorting; phylogenetic network; Bayesian inference; RJMCMC.]

---

The availability of sequence data from multiple loci across the genomes of species and individuals within species is enabling accurate estimates of gene and species evolutionary histories, as well as parameters such as divergence times and ancestral population sizes (Rannala and Yang 2003). Several statistical methods have been developed for obtaining such estimates (Bouckaert *et al.* 2014; Edwards *et al.* 2007; Heled and Drummond 2010; Rannala and Yang 2003). All these methods employ the *multispecies coalescent* (Degnan and Rosenberg 2009) as the stochastic process that captures the relationship between species trees and gene genealogies.

As evidence of hybridization (admixture between different populations of the same species or across different species) continues to accumulate (Arnold 1997; Barton 2001; Gogarten *et al.* 2002; Koonin *et al.* 2001; Mallet 2005, 2007; Rieseberg 1997), there is a pressing need for statistical methods that infer species phylogenies, gene trees, and their associated parameters in the presence of hybridization. We recently introduced for this purpose the *multispecies network coalescent* (MSNC) along with a maximum likelihood search heuristic (Yu *et al.* 2014) and a Bayesian sampling technique (Wen *et al.* 2016a). However, these methods use gene tree estimates as input. Using these estimates, instead of using the sequence data directly, has at least three drawbacks. First, the sequence data allows for learning more about the model than gene tree estimates (Rannala and Yang 2003). Second, gene tree estimates could well include erroneous information, resulting in wrong inferences (DeGiorgio and Degnan 2014; Wen *et al.* 2016a). Third, co-estimating the species phylogeny and gene trees results in better estimates of the gene trees themselves (Bayzid and Warnow 2013; DeGiorgio and Degnan 2014).

We report here on a Bayesian method for co-estimating species (or, population) phylogenies and gene trees along with parameters such as ancestral population sizes and divergence times using DNA sequence alignments from multiple independent loci. Our method utilizes a two-step generative process (Fig. 1) that links, via latent variables that correspond to local gene genealogies, the sequences of multiple, unlinked loci from across a set of genomes to the phylogenetic network (Nakhleh 2010a) that models the evolution of the genomes themselves.

**FIGURE 1.**
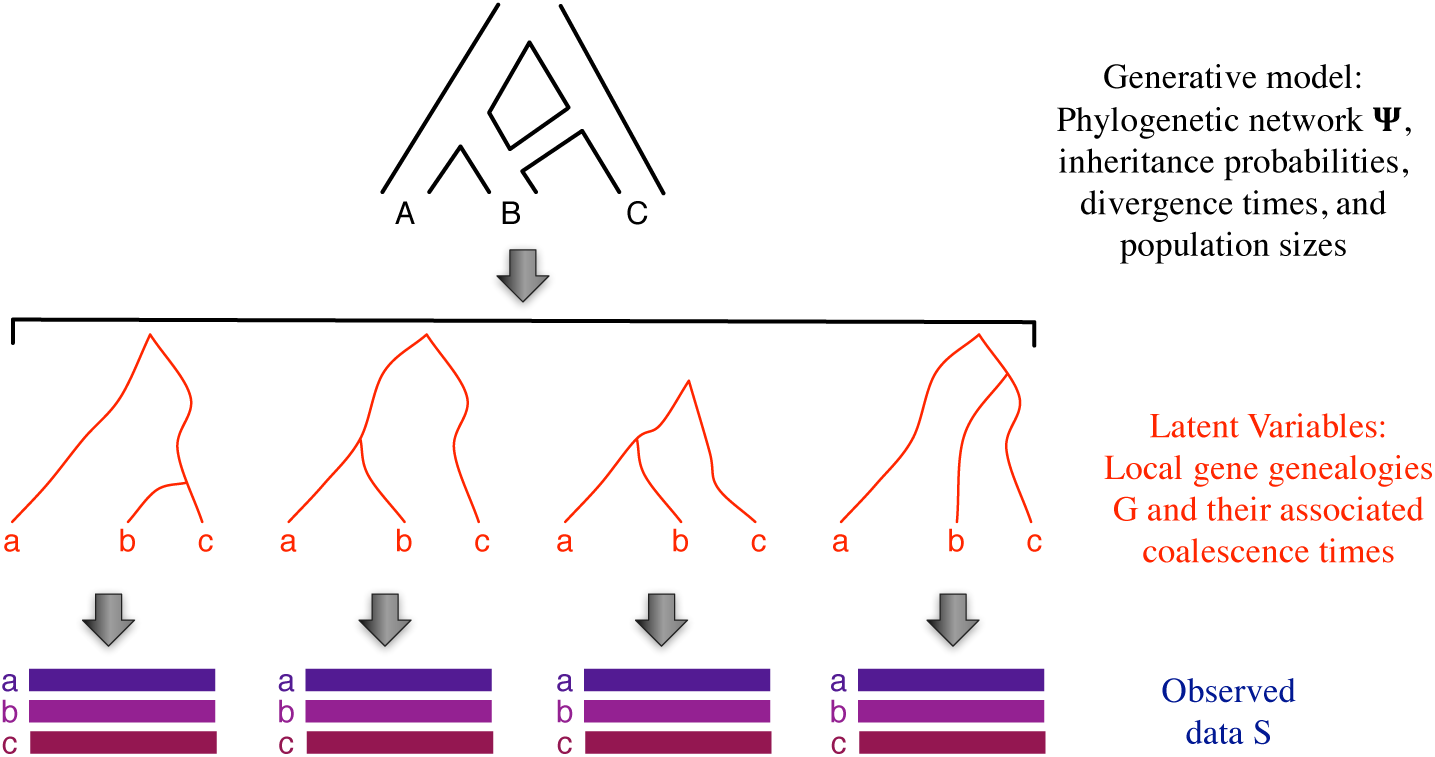
From a phylogenetic network to multi-locus sequences via latent gene genealogies. The multispecies network coalescent (Yu et *al.* 2014) is a stochastic process that defines a probability distribution on gene genealogies along with their coalescent times. The parameters of the process consist of a phylogenetic network topology, inheritance probabilities, divergence times, and population sizes. Each gene genealogy, when coupled with model of sequence evolution, defines a probability distribution on sequence alignments.

Our method consists of a reversible-jump Markov chain Monte Carlo (RJMCMC) sampler of the posterior distribution of this generative process. In particular, our method co-estimates, in the form of posterior distribution samples, the phylogenetic network and its associated parameters for the genomes as well as the local genealogies for the individual loci. We demonstrate the performance of our method on simulated data. Furthermore, we analyze three biological data sets, and discuss the insights afforded by our method. In particular, we find that methods that do not account, wrongly, for admixture in the data tend to underestimate divergence times of the species or populations and overestimate the coalescent times of individual gene genealogies. Our method, on the other hand, estimates both the divergence times and coalescent times with high accuracy. Furthermore, we demonstrate that coalescent times are much more accurately estimated when the estimation is done simultaneously with the phylogenetic network than when the estimation is done in isolation.

An important contribution of this manuscript is also to study the performance of the MSNC on data generated under gene flow scenarios. In particular, the population genetics community has developed models of reticulate evolution (i.e., admixture) at the population level. An important question is: How do phylogenetics network methods perform on data generated under such scenarios? To answer this question, it is important to highlight the difference in abstraction employed in the MSNC model as opposed to a gene flow model. It turns out that this difference was well articulated in (Long 1991), where two models of admixture were presented: the intermixture model and the gene flow model (Figure 2). The MSNC employs the intermixture model, whereas the population genetics community mostly uses the gene flow model (Gronau *et al.* 2011; Hey and Nielsen 2004, 2007; Leaché *et al.* 2013; Slatkin and Maddison 1989; Strasburg and Rieseberg 2010; Whitlock and Mccauley 1999). Note that the intermixture model also underlies the admixture graph model of (Pickrell and Pritchard 2012; Reich *et al.* 2009) where *γ* is the admixture proportion. In the admixture graph model, the branch lengths correspond to genetic drift values that measure variation in allele frequency corresponding to random sampling of alleles from generation to generation in a finite-size population.

**FIGURE 2.**
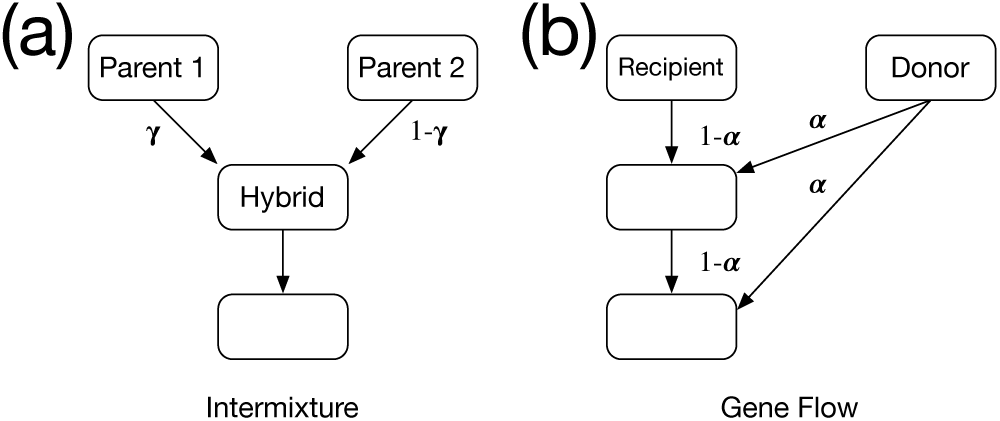
Two admixture models for a hybrid population (Long 1991). (a) The hybrid population is formed by a single intermixture event between two parental populations, where γ is the inheritance probability measuring the proportion of the parental populations. (b) The hybrid population (recipient) receives gene flow from a donor population, where *α* is the migration rate.

Hudson’s ms program (Hudson 2002) allows for generating data under each of the two admixture models—intermixture and gene flow. In this paper, we generate data under both models and study the performance of inference under the MSNC in both cases.

For an empirical data set, we analyzed the yeast data set of (Rokas *et al.,* 2003), which consists of 106 loci from seven Saccharomyces species, and contrasted our results to those obtained from the method of (Wen *et al.,* 2016a) on gene tree estimates.

Finally, as the model underlying our method extends the multispecies coalescent to cases that include admixture, our method is applicable to data from different sub-populations, not only different species, and to data where more than one individual per species or sub-population is sampled. The method is implemented and publicly available in the PhyloNet software package (Than *et al.* 2008).

## METHODS

### 0.1 Phylogenetic networks and their parameters

A *phylogenetic 𝒳 -network,* or *𝒳*-network for short, Ψ, is a directed, acyclic graph (DAG) with *V*(Ψ) = {s,r}U*V*_*L*_U*V*_*T*_U*V*_*N*_, where

- *indeg(s)=0* and *outdeg(s) = 1 (s* is a special node, that is the parent of the root node, r);
- *indeg(r) = 1* and *outdeg(r)* = 2 (r is the *root* of Ψ);
- ∀ *v* ∈ V_*L*_, *indeg(v)* = 1 and *outdeg(v)=0 (V*_*L*_ are the *external tree nodes,* or *leaves,* of Ψ);
- ∀ *v* ∈ *V*_*T*_, *indeg(v)* = 1 and *outdeg(v)* ≥ 2 (*V*_*T*_ are the *internal tree nodes* of Ψ); and,
- ∀ *v* ∈ *V*_*N*_, *indeg(v)* = 2 and *outdeg(v)* = 1 (*V*_*N*_ are the *reticulation nodes* of Ψ).

The network’s edges, E(Ψ)⊆*V* × *V*, consist of *reticulation edges*, whose heads are reticulation nodes, *tree edges*, whose heads are tree nodes, and special edge (*s*,*r*) ∈ *E*. Furthermore, 𝓁: *V*_*L*_ → 𝒳 is the *leaf-labeling* function, which is a bijection from *V*_*L*_ to 𝒳. Each node in *V*(Ψ) has a species divergence time parameter and each edge in *E*(Ψ) has an associated population size parameter. The edge *er*(Ψ) = (*s,r*) is infinite in length so that all lineages that enter it coalesce on it eventually. Finally, for every pair of reticulation edges *e*_1_ and *e*_2_ that share the same reticulation node, we associate an inheritance probability, *γ*, such that *γ*_*e*1_*,γ*_*e2*_ ∈ [0,1] with *γ*_*e*1_ + *γ*_*e2*_ = 1. We denote by Γ the vector of inheritance probabilities corresponding to all the reticulation nodes in the phylogenetic network (for each reticulation node, Γ has the value for one of the two incoming edges only).

Given a phylogenetic network Ψ, we use the following notation:

- *Ψ*_*top*_: The leaf-labeled topology of Ψ; that is, the pair (*V,E*) along with the leaf-labeling *l*.
- Ψ_*ret*_: The number of reticulation nodes in Ψ. Ψ_*ret*_ = 0 when Ψ is a phylogenetic tree.
- Ψ_*τ*_ : The species divergence time parameters of Ψ. Ψ_*τ*_. ∈ (ℝ ^+^)^| *V*(Ψ)|^.
- *Ψ*_*θ*_: The population size parameters of Ψ. Ψ_*θ*_ ∈ (ℝ^+^)^|*E*(Ψ)|^

We use Ψ to refer to the topology, species divergence times and population size parameters of the phylogenetic network.

It is often the case that divergence times associated with nodes in the phylogenetic network are measured in units of years, generations, or coalescent units. On the other hand, branch lengths in gene trees are often in units of expected number of mutations per site. We convert estimates back and forth between units as follows:

- Given divergence time in units of expected number of mutations per site *τ*, mutation rate per site per generation *μ* and the number of generations per year *g*, *τ/μg* represents divergence times in units of years.
- Given population size parameter in units of population mutation rate per site *θ*, *2τ/θ* represents divergence times in coalescent units.

*Bayesian Formulation and Inference*

The data in our case is a set *𝒮* = {S_1_,…, *S*_*m*_*}* where *S*_*i*_ is a DNA sequence alignment from locus *i* (the bottom part in Fig. 1). A major assumption is that there is no recombination within any of the *m* loci, yet there is free recombination between loci. The model ℳ consists of a phylogenetic network Ψ (the topology, divergence times, and population sizes) and a vector of inheritance probabilities Γ (the top part in Fig. 1).

The posterior distribution of the model is given by

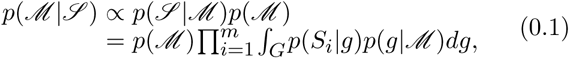

where the integration is taken over all possible gene trees (the middle part in Fig. 1). The term *p(S*_*i*_ |*g*) gives the gene tree likelihood, which is computed using Felsenstein’s algorithm (Felsenstein 1981) assuming a model of sequence evolution, and *p*(*g*|ℳ) is the probability density function for the gene trees, which was derived for the cases of species tree and species network in (Rannala and Yang 2003) and (Yu *et al.* 2014), respectively.

The integration in Eq. (0.1) is computationally infeasible except for very small data sets. Furthermore, in many analyses, the gene trees for the individual loci are themselves a quantity of interest. Therefore, to obtain gene trees, we sample from the posterior distribution as given by

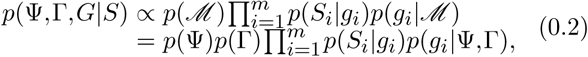

where *G* = (*g*_1_*,…,g*_m_) is a vector of gene trees, one for each of the *m* loci. This co-estimation approach is adopted by the two popular Bayesian methods *BEAST (Heled and Drummond 2010) and BEST (Liu 2008), both of which co-estimate species trees (hybridization is not accounted for) and gene trees.

*The Likelihood Function*

Felsenstein (Felsenstein 1981) introduced a pruning algorithm that efficiently calculates the likelihood of gene tree *g* and DNA evolution model parameters Φ as

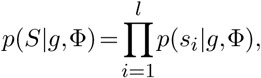

where *s*_*i*_ is i-th site in S, *l* is the sequence length, and

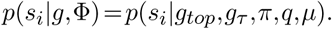

Here, *g*_*top*_ is the tree topology, *g*_*τ*_ is the divergence times of the gene tree, *π* = {*π*_*A*_*,π*_*T*_*,π*_*C*_*,πG*} is a vector of equilibrium frequencies of the four nucleotides, *q* = {*qAT, qAC ,qAG,qTC,qTG,qCG*} is a vector of substitution rates between pairs of nucleotides, and μ is the mutation rate. Over a branch *j* whose length (in expected number of mutations per site) is *t*_*j*_, the transition probability is calculated as *e*^μ *qt_j_*^. In the implementation, we use the BEAGLE library (Ayres *et al.* 2011) for more efficient implementation of Felsenstein’s algorithm.

Yu *et al.* (Yu *et al.* 2012, 2013a, 2014) fully derived the mass and density functions of gene trees under the multispecies network coalescence, where the lengths of a phylogenetic network’s branches are given in coalescent units. Here, we derive the probability density function (pdf) of gene trees for a phylogenetic network given by its topology, divergence/migration times and population size parameters following (Rannala and Yang 2003; Yu *et al.* 2014). Coalescence times in the (sampled) gene trees posit temporal constraints on the divergence and migration times of the phylogenetic network.

We use τΨ(v) to denote the divergence time of node *v* in phylogeny Ψ (tree or network). Given a gene tree g whose coalescence times are given by τ' and a phylogenetic network Ψ whose divergence times are given by τ, we define a coalescent history with respect to times to be a function h: V(g) →E(Ψ), such that the following condition holds:

- if (*x,y*) ∈ *E*(Ψ) and 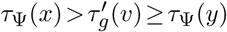, then h(v) = (*x, y*).
- if *r* is the root of Ψ and 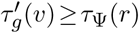, then *h*(*v*) = er(Ψ).

The quantity 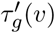 indicates at which point of branch (*x,y*) coalescent event v happens. We denote the set of coalescent histories with respect to coalescence times for gene tree g and phylogenetic network Ψ by *H*_Ψ_ (g).

Given a phylogenetic network Ψ, the pdf of the gene tree random variable is given by

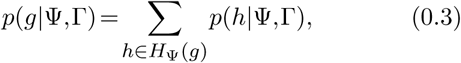

where *p*(*h*|Ψ, Γ) gives the pdf of the coalescent history (with respect to divergence times) random variable.

Consider gene tree *g* for locus *j* and an arbitrary *h* ∈ *H*_Ψ_(g). For an edge b = (*x,y*) ∈ E(Ψ), we define *T*_*b*_(*h*) to be a vector of the elements in the set {τ_*g*_ (*w*): *w* ∈ h^-1^(b)}U{τ_Ψ_(y)} in increasing order. We denote by *T*_b_(h)[*i*] the *i*-th element of the vector. Furthermore, we denote by *u*_*b*_(*h*) the number of gene lineages entering edge *b* and v_*b*_(*h*) the number of gene lineages leaving edge *b* under *h*. Then we have

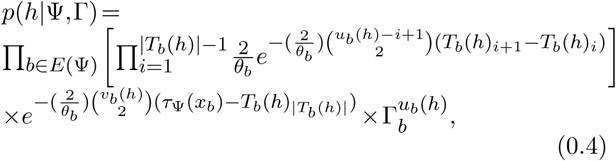

where *x*_*b*_ is the source node of edge *b*, θ_*b*_ = 4N_b_*μ* and *N*_*b*_ is the population size corresponding to branch *b*, μ is the mutation rate per-site per-generation, and Γ_*b*_ is the inheritance probability associated with branch *b*.

*Prior Distributions*

We extended the prior of phylogenetic network composed of topology and branch lengths in (Wen *et al.* 2016a) to phylogenetic networks composed of topology, divergence times and population sizes, as given by Eq. (0.5),

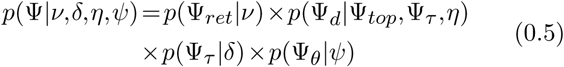

where *p*(Ψ_*ret*_|*v*), the prior on the number of reticulation nodes, and *p*(Ψ_d_|*Ψ*_*top*_,*Ψ*_*τ*_,η), the prior on the diameters of reticulation nodes, were defined in (Wen *et al.* 2016a).

It is important to note here that if *Ψ*_*top*_ does not follow the phylogenetic network definition, then *p*(Ψ|*v*, *δ*, *η*, *ψ*) = 0. This is crucial since, in the MCMC kernels we describe below, we allow the moves to produce directed graphs that slightly deviate from the definition; in this case, having the prior be 0 guarantees that the proposal is rejected. Using the strategy, rather than defining only “legal” moves simplifies the calculation of the Hastings ratios. See more details below.

Rannala and Yang used independent Gamma distributions for time intervals (branch lengths) instead of divergence times. However, in the absence of any information on the number of edges of the species network as well as the time intervals, it is computationally intensive to infer the hyperparameters of independent Gamma distributions. Currently, we use a uniform distribution (as in BEST (Liu 2008)).

We assume one population size per edge, including the edge above the root. Population size parameters are Gamma distributed, *θ*_*b*_ ∼ Γ (2, *ψ*), with a mean 2*ψ* and a shape parameter of 2. In the absence of any information on the population size, we use the noninformative prior *P*_*ψ*_ (*x*) = 1/*x* for hyperparameter *ψ* (Heled and Drummond 2010). The number of elements in θ is |E(*Ψ*)| + 1. To simplify inference, our implementation also supports a constant population size across all branches, in which case θ contains only one element.

For the prior on the inheritance probabilities, we use Γ_*b*_∼Beta(*α*, *β*). Unless there is some specific knowledge on the inheritance probabilities, a uniform prior on [0,1] is adopted by setting *α* = *β* =1. If the amount of introgressed genomic data is suspected to be small in the genome, the hyper-parameters *α* and *β* can be appropriately set to bias the inheritance probabilities to values close to 0 and 1 (a U-shaped distribution).

### The RJMCMC Sampler

As computing the posterior distribution given by Eq. (0.2) is computationally intractable, we implement a Markov chain Monte Carlo (MCMC) sampling procedure based on the Metropolis-Hastings algorithm. In each iteration of the sampling, a new state (Ψ′, Γ′,*G*′) is proposed and either accepted or rejected based on the Metropolis-Hastings ratio *r* that is composed of the likelihood, prior, and Hastings ratios. When the proposal changes the dimensionality of the sample by adding a new reticulation to or removing an existing reticulation from the phylogenetic network, the absolute value of the determinant of the Jacobian matrix is also taken into account, which results in a reversible-jump MCMC, ors RJMCMC (Green 1995, 2003).

Our sampling algorithm employs three categories) of moves: One for sampling the phylogenetic network and its parameters (divergence times and population mutation rates), one for sampling the inheritance probabilities, and one for sampling the gene trees (topologies and coalescence times). To propose a new state of the Markov chain, one element from (Ψ,γ1,…,γΨ_ret_,g1*,…,g*_*m*_) is selected at random, then a move from the corresponding category is applied. The workflow, design and full derivation of the Hastings ratios of the moves are given in Supplementary Materials.

We implemented our method in PhyloNet (Than *et al.* 2008), a publicly available, open-source software package for phylogenetic network inference and analysis.

## RESULTS

### Our Method and *BEAST Perform Similarly in Cases of No Reticulation

*BEAST (Heled and Drummond 2010) is the most commonly used software tool for Bayesian inference of species trees from multi-locus data. In our first experiment, we set out to study how our method performs compared to this well-established software tool on simulated data whose evolutionary history is treelike. To accomplish this task, we used the phylogenetic tree shown in Fig. 3 as the model species phylogeny. Using the program ms (Hudson 2002), we simulated 20 data sets each consisting of 10 conditionally independent gene trees with the command

**FIGURE 3.**
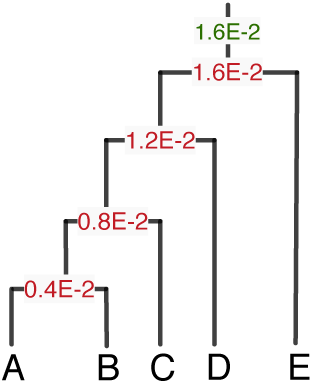
A model species tree used to generate multi-locus data sets. The divergence times in units of expected number of mutations per site and the population size parameter in units of population mutation rate per site are marked in red and green, respectively. The population mutation rate was assumed to be constant across all branches of the tree.

ms 5 10 -T -I 5 1 1 1 1 1 -ej 0.25 3 2 -ej 0.5 4 2 -ej 0.75 5

2 -ej 1.0 2 1

We then used the program Seq-gen (Rambaut and. Grassly 1997) to simulate the evolution of 1000-site_(_ sequences under the Jukes-Cantor model of evolution (Jukes and Cantor 1969) with the command

seq-gen -m HKY -l 1000 -s 0.008

For each of the 20 10-locus data sets, we ran two: MCMC chains, each with 5 × 10^5^ iterations and 5 x: 10^4^ burn-in, using our method as well as *BEAST. One sample was collected from every 500 iterations, resulting in a 900 collected samples per data set and a total of 18,000 collected samples from all 20 data sets. In comparing the two tools, we used all 18,000 collected samples to evaluate the estimates obtained for the various parameters of interest: population size parameter, divergence times, and the topology of the inferred species phylogeny.

Both our method and *BEAST inferred exactly the same 95% credible set, which consists of the six topologies shown in Fig. 4. Our method sampled the true phylogeny with higher frequency than *BEAST.

**FIGURE 4.**
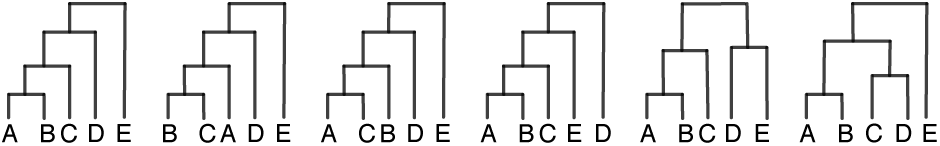
The trees that constitute the 95% credible set of each of our method and *BEAST. The proportions of these trees from left to right as sampled by our method were 77.7%, 5.7%, 5.0%, 3.0%, 3.0%, and 2.8%, respectively, and as sampled by *BEAST were 70.7%, 6.0%, 6.7%, 4.7%, 4.5%, and 3.6%, respectively.

Fig. 5 shows histograms of the estimates obtained for the divergence times at each node of the maximum a posteriori (MAP) species tree estimate of our method and *BEAST, which was identical in both cases to the true species tree. The histograms of both methods are very similar. In fact, the histograms obtained by our method have peaks that are closer to the true divergence time values than those obtained by *BEAST.

**FIGURE 5.**
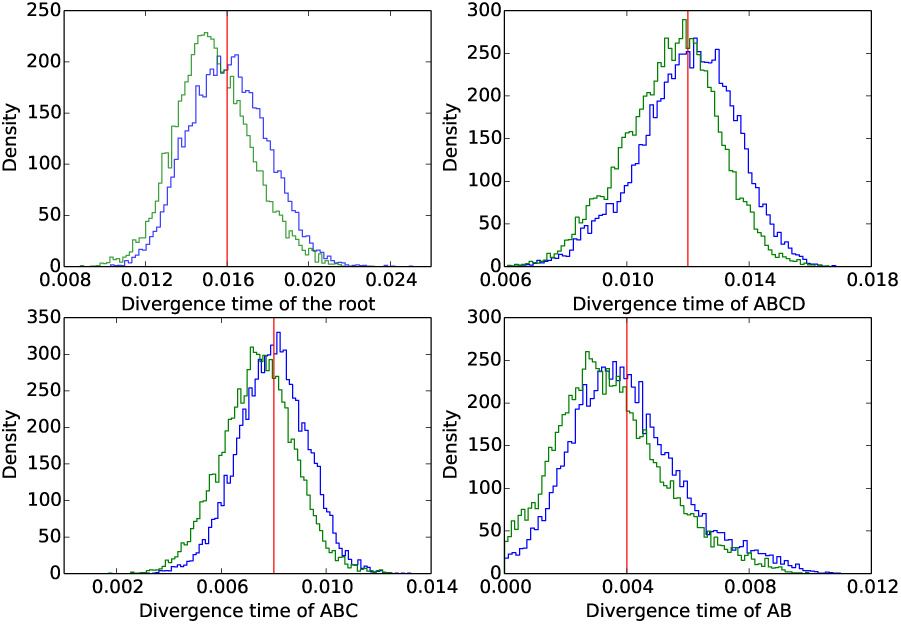
Histograms of divergence times of each node of the true species phylogeny as estimated by our method (blue) and *BEAST (green). The red vertical line indicates the true divergence time.

Fig. 6 shows the histograms of the population mutation rate (one value across all branches of the species tree was assumed) estimated by the two methods. As in the case of divergence time estimates, the two methods obtain similar results in the case of population mutation rate estimates. However, we observe here a histogram of our method with a single peak around the true value, whereas we observe a bimodal histogram obtained by *BEAST.

**FIGURE 6.**
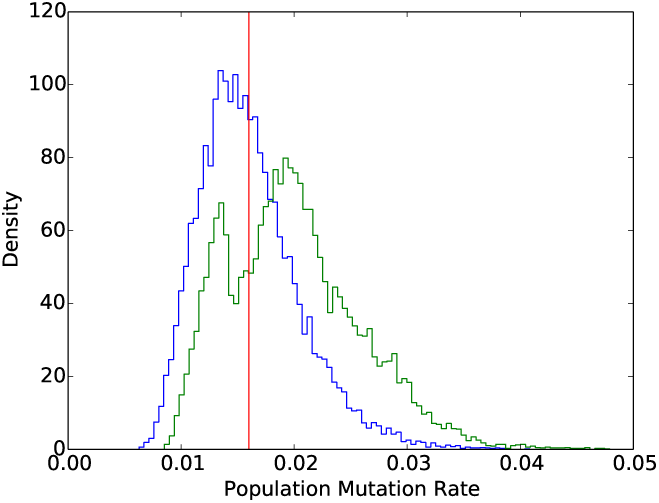
Population mutation rate estimated by our method (blue) and *BEAST (green). The red vertical line indicates the true population mutation rate.

All the results reported above were obtained by running the code on NOTS (Night Owls Time-Sharing Service), which is a batch scheduled High-Throughput Computing (HTC) cluster. We used 2 cores, with two threads per core running at 2.6GHz, and 1G RAM per thread. The runtime for *BEAST is around 28±1 seconds for each data set, while our method takes longer time: 185±7 seconds per data set. This can be explained by the fact that *BEAST has been under continued development for several years now, while our implementation hardly has any optimization components yet.

When we ran *BEAST on multi-locus sequence data simulated under species phylogenies with reticulations, we found that *BEAST overestimated the coalescence times in individual loci and underestimated the divergence times of the species phylogeny. We report these results in Supplementary Materials as *BEAST is not intended for evolutionary analyses with gene flow. Furthermore, there are existing, extensive studies on the impact of gene flow on the inference of species trees (Leache *et al.* 2013; Solis-Lemus *et al.* 2016).

### Our Method Provides Accurate Estimates of the Network and Its Associated Parameters

We used the phylogenetic network shown in Fig. 7 as the model species phylogeny. The scale parameter of the divergence times s was varied to take on values in the set {0.1,0.25,0.5,1.0}. Setting s = 0.1 results in very short branches and, consequently, the hardest data sets on which to estimate parameters. Setting s = 1.0 results in longer branches and higher signal for a more accurate estimate of the parameter values. It is important to note that the topology, reticulation event, divergence times (with s = 1.0) and population size are inspired by the species phylogeny recovered from the Anopheles mosquitoes data set (Fontaine *et al.* 2015; Wen *et al.* 2016b).

For the four settings of s values, 0.1, 0.25, 0.5, and 1.0, we used the program ms (Hudson 2002) to simulate 20 data sets each with 128 gene trees of conditionally independent loci with the four following commands respectively.

**FIGURE 7.**
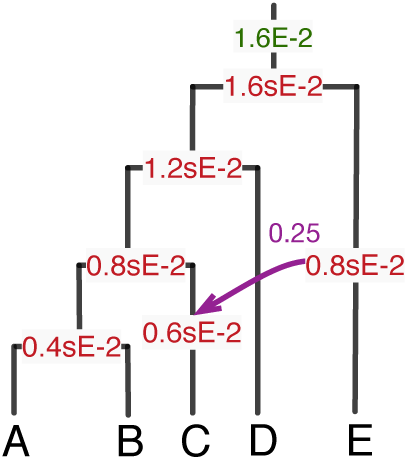
A model phylogenetic network used to generate simulated data. The divergence times in units of expected number of mutations per site, the population size parameter in units of population mutation rate per site, and the inheritance probability are marked in red, green, and purple, respectively. Parameter s is used to scale the divergence times.

- ms 5 128 -T -I 5 1 1 1 1 1 -ej 0.025 4 3 -es 0.0375 1 0.3 -ej 0.05 6 3 -ej 0.05 2 1 -ej 0.075 5 3 -ej 0.1 3 1
- ms 5 128 -T -I 5 1 1 1 1 1 -ej 0.0625 4 3 -es 0.09375 1 0.3 -ej 0.125 6 3 -ej 0.125 2 1 -ej 0.1875 5 3 -ej 0.25 3 1
- ms 5 128 -T -I 5 1 1 1 1 1 -ej 0.125 4 3 -es 0.1875 1 0.3 -ej 0.25 6 3 -ej 0.25 2 1 -ej 0.375 5 3 -ej 0.5 3 1
- ms 5 128 -T -I 5 1 1 1 1 1 -ej 0.25 4 3 -es 0.375 1 0.3 -ej 0.5 6 3 -ej 0.5 2 1 -ej 0.75 5 3 -ej 1.0 3 1

The program Seq-gen (Rambaut and Grassly 1997) was used to generate sequence alignments down the gene trees under the Jukes Cantor model (Jukes and Cantor 1969) with lengths seqLen in {250,500,1000} using the command

### seq-gen -m HKY -l seqLen -s 0.008

To vary the number of loci used in the inference, we produced data sets with 32, 64, and 128 loci by sampling loci without replacement from the full data set of 128 loci. Each of these sequence data sets was then used as input to the inference method.

To assess the signal in the sequence data sets we6 obtained, we quantified the percentage of variable sites for each setting, averaged over all 20 replicates for that setting. The percentages of variable sites in the generated alignments for *s* = 0.1,0.25,0.5,1.0 (varying the sequence length had negligible effect for the same scaling factor s) are ∼0.039±0.02, ∼0.048±0.02, ∼0.061±0.02, and ∼0.088±0.02, respectively.

For each data set, we ran an MCMC chain of 8x 10^6^ iterations with 1 × 10^6^ burn-in. One sample was collected from every 5,000 iterations, resulting in a total of 1,400 collected samples. We summarized the results based on 28,0 samples from 20 replicates for each of the 36 simulation settings (four values of *s*, three sequence lengths, and three numbers of loci). In the boxplots below, the five bars from bottom to top correspond to the minimum, first-, second-, third-quantile, and the maximum, respectively, from the 20 replicates for each setting. In the other figures, the error bars correspond to standard deviations calculated from the 20 replicates for each setting.

In assessing the performance of our method, we evaluated the estimates obtained for the various parameters of interest: divergence times, population mutation rates, the number of reticulations, and the topology of the inferred species phylogeny. Fig. 8 shows the estimates obtained for the divergence time at the root of the network. Three observations are in order.

**FIGURE 8.**
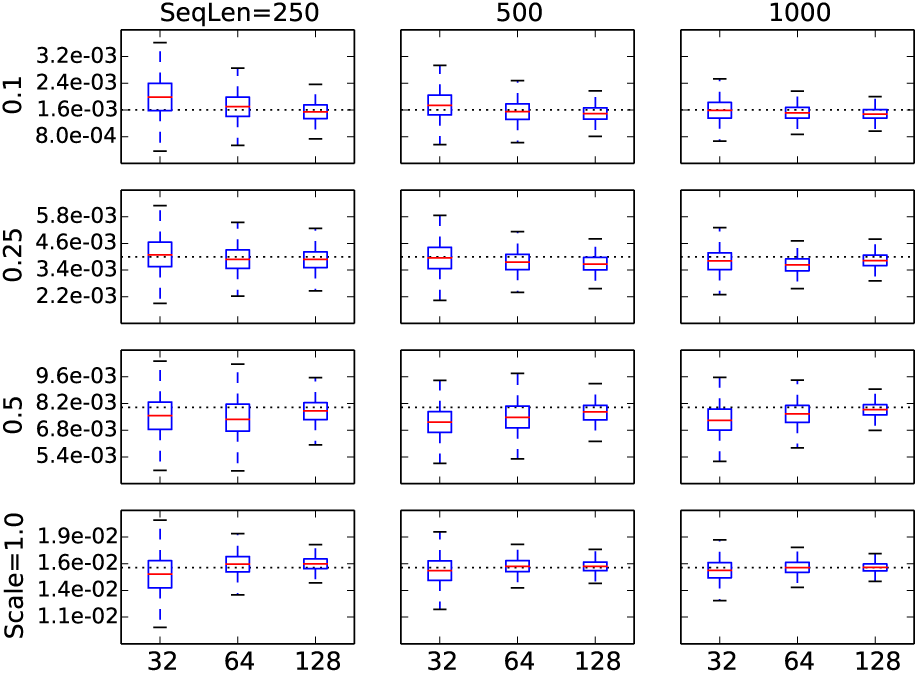
Divergence time estimates at the root under different values of the scaling parameter *s* (different rows), sequence lengths (different columns), and numbers of loci (three values within each panel). The dashed line indicates the true value in the model network.

First, for any combination of sequence length and scaling parameter value, the divergence time estimate converges to the true value as the number of loci increases. Second, for any combination of number of loci and scaling parameter value, the divergence time estimate converges to the true value as the sequence length increases. Third, the estimates are relatively poor only under the extreme settings of scaling parameter value 0.1 and sequence length 250. In this case, the signal in the sequence data is too weak to obtain good estimates. However, it is worth noting that even under this setting, using 128 loci produces a very accurate estimate of the divergence time.

Fig. 9 shows the estimates obtained for the population mutation rate parameter (one value across all branches of the species network was assumed). The results show very similar trends to those obtained for the divergence time estimates, with the main difference being that the estimates now are very accurate even for the hardest of cases: *s* = 0.1 and sequence length 250, regardless of the number of loci used.

**FIGURE 9.**
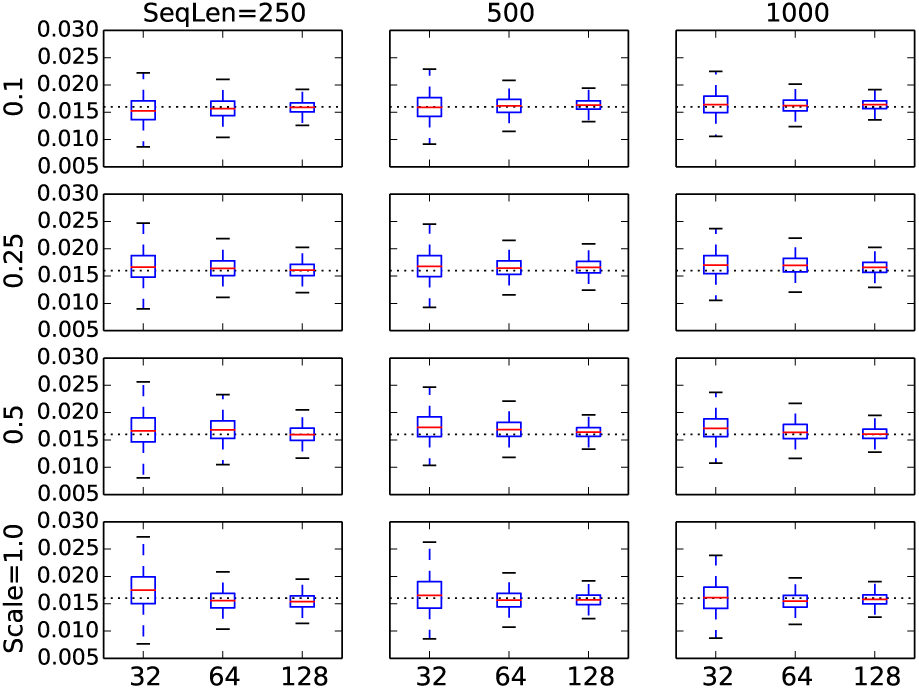
Population mutation rate estimates under different values of the scaling parameter *s* (different rows), sequence lengths (different columns), and numbers of loci (three values within each panel). The dashed line indicates the true value in the model network.

The results are quite different when it comes to estimating the number of reticulations and the topology of the phylogenetic network itself. Fig. 10 shows the estimates of the number of reticulations under different settings. As the figure clearly shows, under the case of extremely short branches (*s* = 0.1), the method recovers a tree; that is, it estimates the number of reticulations to be 0, regardless of the number of loci or sequence length used. Here, the signal is too weak to recover any reticulation. In the case of slightly longer branches (s = 0.25), the estimate of the number of reticulations becomes slightly more accurate when the sequences are long and 128 loci are used. Given the observed trend, the method could recover the true number of reticulations if a thousand or so loci are used. In the case of *s* = 0.5, a fast convergence towards the true number is observed as the number of loci increases. It is worth pointing out that, in the case of *s* = 0.5, increasing the number of loci, even when the sequences are very short, is much more advantageous than increasing the sequence lengths of the individual loci. It is also important to note here that in analyzing biological data sets, one cannot use longer sequences without risking violating the recombination-free loci assumption. In the case of *s* = 1. 0, the method does very well at estimating the number of reticulations. Finally, observe that the method almost never overestimates the number of reticulations on these data sets.

**FIGURE 10.**
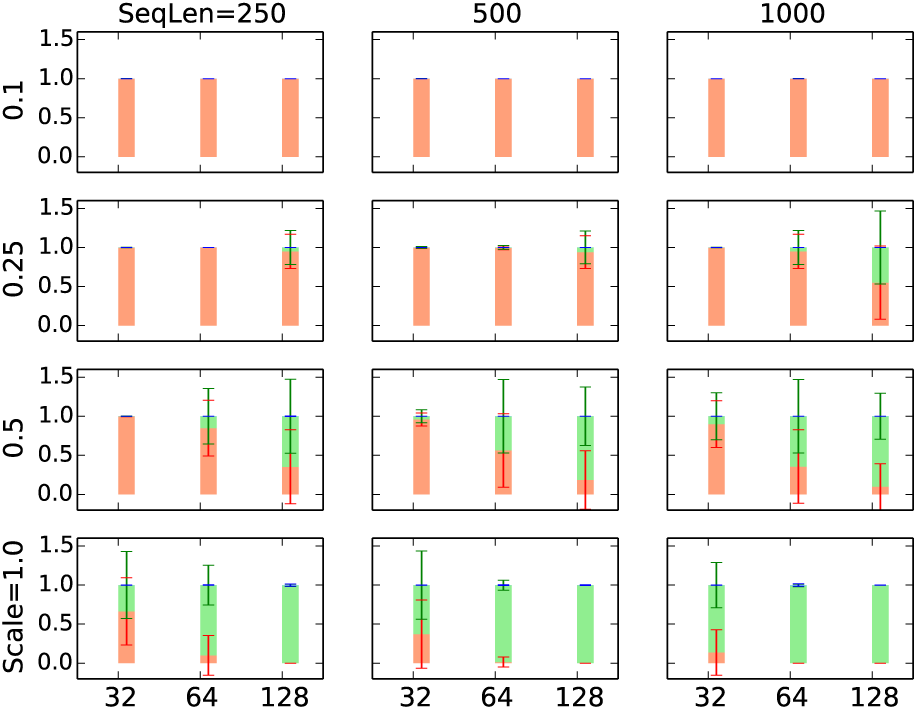
Proportions of trees (red), 1-reticulation networks (green) and 2-reticulations networks (blue) inferred under different simulation conditions. The model network has a single reticulation.

In assessing the quality of the estimated network topology itself, we analyzed the recovered networks in two ways. First, we compared the inferred network to the true network using a topological dissimilarity measure (Nakhleh 2010b). Second, when the method infers a tree, rather than a network, we compared the tree to the “backbone tree” of the true network (the tree resulting from removing the arrow in Fig. 7) using the Robinson-Foulds metric (Robinson and Foulds 1981). The latter comparison allows us to answer the question: When the method estimates the species phylogeny to be a tree, how does this tree compare to the backbone tree of the true network It is important to note, though, that the relationship of a phylogenetic network and its constituent trees can become too complex to be captured by a backbone tree in the presence of incomplete lineage sorting (Zhu *et al.* 2016). Fig. 11 shows the results. The results in terms of the topological difference between the inferred and true networks parallel those that we discussed above in terms of the estimates of the number of reticulations: Poor accuracy and no sign of convergence to the true network in cases of very small values of the scaling parameter, and very good accuracy and fast convergence to accurate estimates in cases of larger values of the scaling parameter. However, the topological difference between the inferred trees (in the cases where trees were inferred) and the backbone tree reveal an important insight: When the method fails to recover the true network, it does a very good job at recovering the backbone tree of the true network.

**FIGURE 11.**
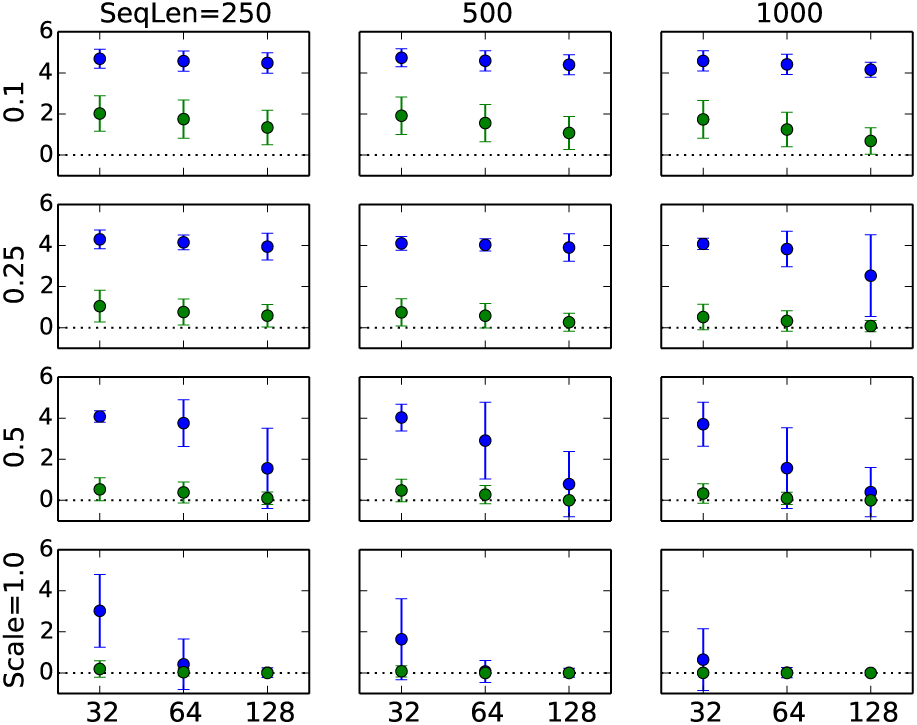
The topological difference between the true and inferred networks in blue and the Robinson-Foulds distance between the inferred tree (if a network is inferred, this case is not included) and the backbone tree of the true network in green.

### Our Method Provides Accurate Estimates of the Gene Trees

Thus far, we have analyzed the accuracy of the inferred networks and their associated parameters. While MCMC methods in this context are deployed to approximate the integration over gene trees in a simulated manner, the methods do provide the sampled gene trees (topologies and coalescence times). The accuracy of those sampled gene trees is important for at least two reasons. First, their accuracy directly impacts and explains the accuracy of the networks. Second, the gene trees themselves are a quantity of interest in many applications.

It is important to note here two relevant studies that have addressed the issue of gene tree accuracy in the context of species tree estimation. First, (Bayzid and Warnow 2013) showed that *BEAST yields more accurate gene trees than would be estimated by RAxML, attributing the higher accuracy to the co-estimation nature of the former method. Second, (DeGiorgio and Degnan 2014) found that methods for estimating gene trees do a better job at estimating the topologies than the coalescence times and that this leads to more accurate species tree estimates when using gene tree topologies alone as opposed to using coalescence times as well. While both studies were conducted in the context of species trees, our goal here is not to reproduce these extensive studies in the context of phylogenetic networks, but rather to demonstrate that the main conclusions still hold even when the species phylogeny is reticulate.

In Fig. 12 we report the Robinson-Foulds distances between the true gene tree topologies and those sampled by our method, as well as the distance between the true gene tree topologies and those estimated by RAxML. The results demonstrate that the co-estimated gene tree topologies are, on average, slightly closer to the true gene tree topologies than those estimated in a standalone manner using RAxML. Nonetheless, it is worth point out that the error bars of our method are smaller than those pertaining to the RAxML gene trees. Both methods obtained improved accuracy as the sequence length increased.

As the results in the next section show, the networks inferred from sequences directly are more accurate than those inferred from gene tree estimates. The question is: What is causing this difference if the gene tree topologies

**FIGURE 12.**
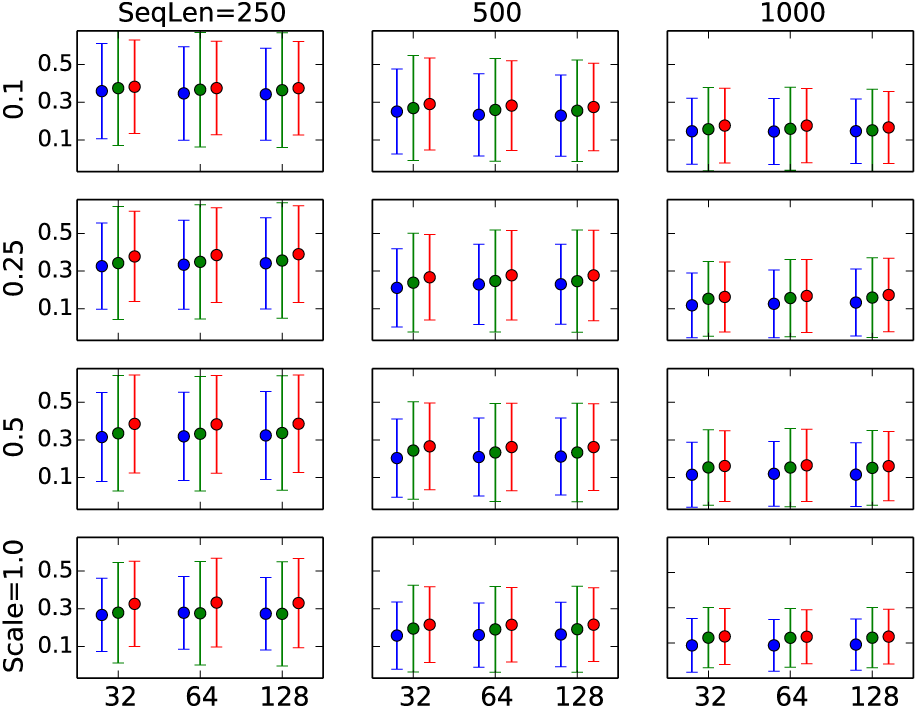
The Robinson-Foulds distances between the true gene tree topologies and those estimated by our method in blue, between the true gene tree topologies and those estimated by RAxML in green, and between the gene tree topologies estimated by our method and those estimated by RAxML in red.

As the results in the next section show, the networks inferred from sequences directly are more accurate than those inferred from gene tree estimates. The question is: What is causing this difference if the gene tree topologies estimated by both our method and RAxML are not that different? One interesting observation we make is that while both our method and RAxML infer gene tree topologies that, on average, are of equal distance from the true gene tree topologies, the two methods return different trees, as shown in Fig. 12. That is, under the Robinson-Foulds distance, both methods infer gene trees whose topologies could be considered to be, roughly, equally good. However, the topologies are not the same. This difference could explain, at least in part, the increased accuracy of the networks and their associated parameters when inferred from sequences as opposed to gene tree estimates.

To further investigae this question, we turned our attention to the accuracy of the coalescence times estimated by our method. Fig. 13 shows the Normalized Rooted Branch Score (NRBS) (Heled and Drummond, 2010) between the gene trees estimated by our method and the true gene trees. This measure takes into account the branch lengths of the gene trees and not only the topologies. These results clearly show that, except for the hardest case of 0.1 scaling factor, the method performs very well in terms of estimating the coalescence times, not only in terms of the mean value but also in terms of the very small standard deviations.

It is important to comment on a seeming discrepancy between Fig. 12 and Fig. 13. For example, in the case of scaling factor 1 . 0, Fig 12 shows a Robinson-Foulds distance of 0.3, yet Fig. 13 shows an NRBS value close to 0. Given that the number of taxa is 5, a Robinson-Foulds value of 0.3 amounts, roughly, to a single incorrect branch in the gene tree. However, while the true and estimated gene tree differ by one branch, the difference in coalescence times between the two trees could be negligible, which explains the small NRBS values.

**FIGURE 13.**
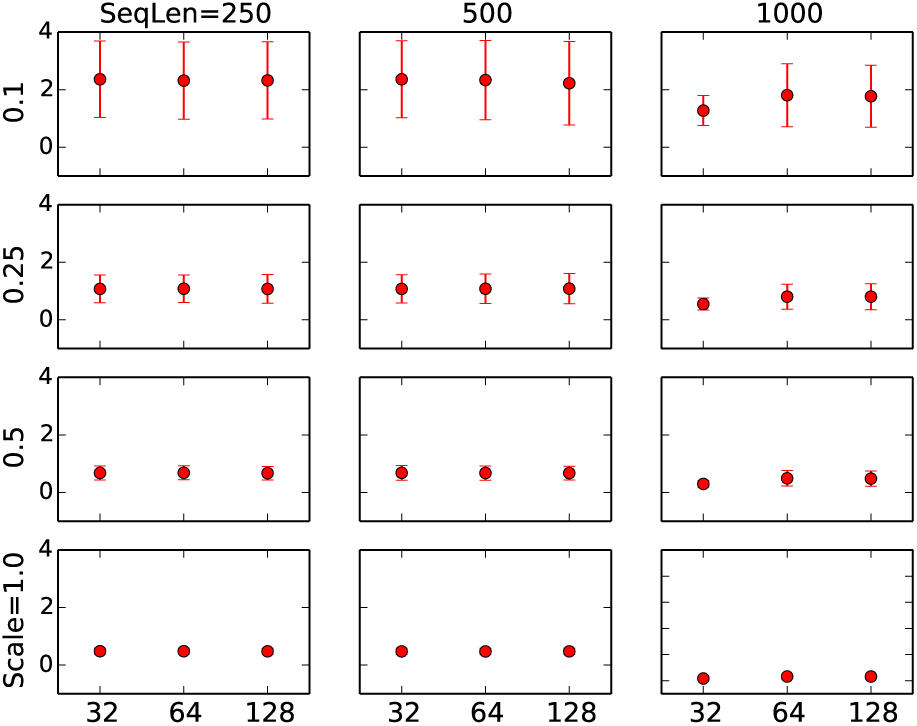
The Normalized Rooted Branch Score (NRBS) (Heled and Drummond, 2010) between the true gene trees and those estimated by our method. The branch lengths are scaled in coalescent units and divided by their corresponding scale parameter 0.1, 0.25, 0.5, 1.0 for better comparison.

Next we show the effect of errors in gene tree estimates on the accuracy of and data requirement for accuracy phylogenetic network estimates.

### Inference from Gene Tree Estimates Requires More Data Than Inference from Sequences

We also set out to compare the performance of our method to that of the method we developed earlier for Bayesian inference of phylogenetic networks from gene tree data (Wen *et al.* 2016a). This method is also implemented in PhyloNet (Than *et al.* 2008) and executed via the command MCMC_GT. The goal here is to assess the gains one obtains by using the sequence data directly rather than first estimating gene trees and then using those as the data for species phylogeny inference.

For the purpose of this experiment we used the subset of the data sets described above and simulated on the phylogenetic network of Fig. 7 under the settings of *s* = 1.0, sequence length 250, and 32, 64, and 128 loci. When using the method of (Wen *et al.* 2016a) we ran it once on the true gene trees and again using the gene tree estimates obtained by RAxML (Stamatakis 2014).

We ran the method of (Wen *et al.* 2016a) for 1,100,000 iterations with 100,000 burn-in and sampled every 1,000 iterations. The top five topologies sampled are shown in Fig. 14 (they were the same top topologies when either the true gene trees or gene tree estimates were used).

**FIGURE 14.**
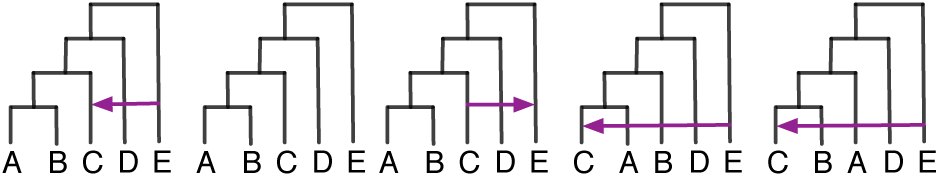
The top five topologies sampled using the method (Wen *et al.,* 2016a) on the true gene trees, as well as the gene tree estimates. The leftmost topology is the true network topology and the second from left is the backbone tree of the true network topology. See the main text for details on the 95% credible sets in terms of these five topologies for the different data sets used.

When using the true gene tree topologies as input data the results were as follows:

- For the 32-locus data set, the 95% credible set contains 16.4% the true network, 59.6% the backbone tree, 12.5% other 1-reticulation networks, and 11.5% other trees.
- For the 64-locus data set, the 95% credible set contains 66.0% the true network, 27.1% the backbone tree, and 3.8% the 1-reticulation network resulting for the backbone tree with reticulation edge *C*→*E* (the network in the middle of Fig. 14).
- For the 128-locus data set, the 95% credible set contains 91.7% the true network, and 4.4% the backbone tree.

When using the gene tree topology estimates as input data, the results were as follows:

- For the 32-locus data set, the 95% credible set contains 6.1% the true network, 47.3% the backbone tree, 14.1% other 1-reticulation networks, and 32.5% other trees.
- For the 64-locus data set, the 95% credible set contains 24.7% the true network, 40.5% the backbone tree, and 8.6% the 1-reticulation network resulting for the backbone tree with reticulation edge *C*→E, 18.4% other 1-reticulation networks, and 7.8% other trees.
- For the 128-locus data set, the 95% credible set contains 49.9% the true network, 19.1 the 1 reticulation network resulting for the backbone tree with reticulation edge *C*→*E*, 5.7% the backbone tree, and 35.2% other 1-reticulation networks.

More comprehensively, Fig. 15 shows the proportions of 0-(tree), 1-, and 2-reticulation networks in the 95% credible sets on each of the data sets when different numbers of loci are used and when the method of (Wen *et al.* 2016a) is run on true and estimated gene tree topologies.

**FIGURE 15.**
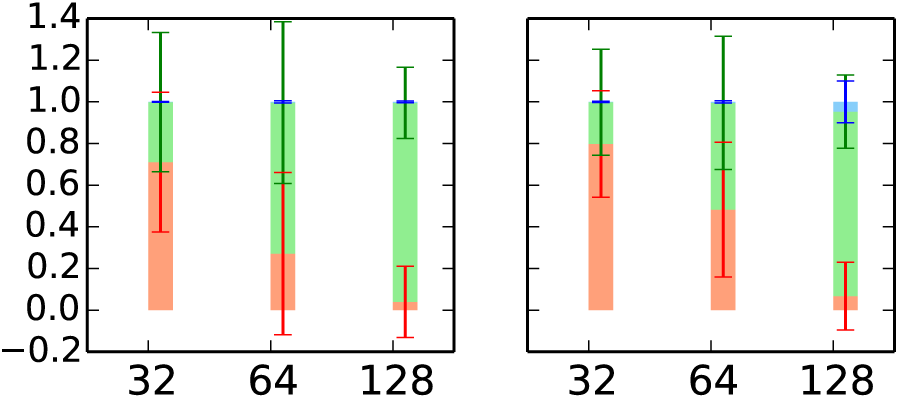
Proportions of trees (red), 1-reticulation networks (green) and 2-reticulations networks (blue) in the 95% credible sets sampled by the method of (Wen *et al.* 2016a) on data sets with 32, 64, and 128 loci. Left: the true gene tree topologies are used as the input data. Right: the gene tree estimates (using RAxML) are used as the input data.

We also assessed the quality of the inferred network/tree topologies by comparing them to the true network using the topological dissimilarity measure (Nakhleh 2010b). When the method infers a tree, rather than a network, we compared the tree to the backbone tree of the true network using the Robinson-Foulds metric (Robinson and Foulds 1981). The results are in Fig. 16.

**FIGURE 16.**
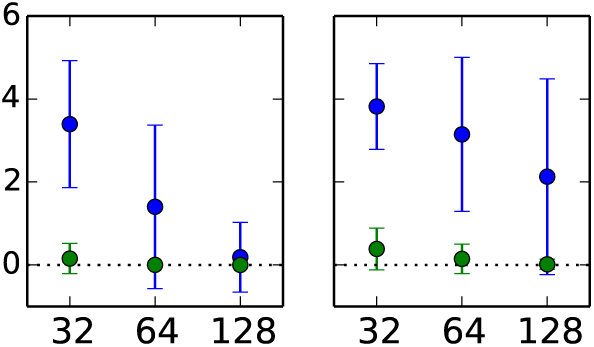
The topological difference between the true and inferred networks in blue and the Robinson-Foulds distance between the inferred tree (if a network is inferred, this case is not included) and the backbone tree of the true network. Left: the true gene tree topologies are used as the input data. Right: the gene tree estimates (using RAxML) are used as the input data.

Clearly, the results indicate the method’s performance in terms of phylogenetic inference improves as the number of loci increases, and, unsurprisingly, the method has a much better performance when the true gene trees are used as input. However, for empirical data sets, the “true” gene trees are never known, and their estimates must be used for methods that utilize gene trees as data. Contrast these results to those obtained by our method When it is run on the sequence data as input (bottom left panel in Fig. 11). Estimation from sequence data outperforms inference from gene trees, even when using the true gene tree topologies. This is mainly due to the fact that the gene tree topology does not capture all the information that the sequence data do. In particular, we observe that inference from sequence data requires a much smaller number of loci than that required to achieve a similar accuracy when making inferences from gene tree topology estimates.

### Intermixture vs Gene Flow: Comparing the Method’s Performance on Data under Both Models

As we discussed above and illustrated in Fig. 2, intermixture and gene flow provide two different abstract models of reticulation. Furthermore, the program ms (Hudson 2002) allows for generating data under both models. While the MSNC is based on an intermixture model, we study here how it performs on data simulated under a gene flow model. We set up the experiment so that data are generated under the same phylogenetic networks and their parameters, yet under the scenarios of intermixture and gene flow separately. Furthermore, in this part, we assess the performance when multiple reticulation events occur between the same pair of species—a very realistic scenario in practice. Fig. 17 shows the six phylogenetic networks we used to generate data.

**FIGURE 17.**
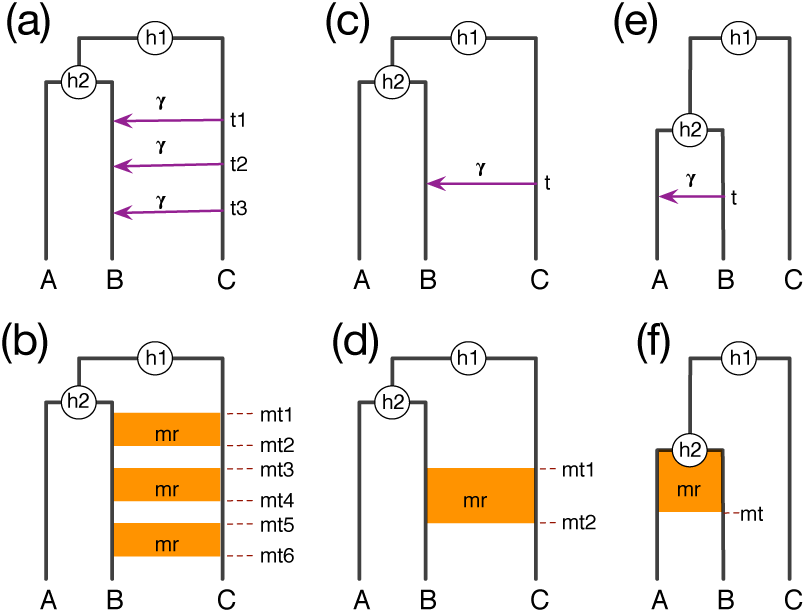
True phylogenetic histories with intermixture and gene flow models. Recurrent reticulations between non-sister taxa (a,b), a single reticulation between non-sister taxa (c,d), and a single reticulation between sister taxa (e,f) are captured under both the intermixture model (top) and gene flow model (bottom). Parameters h1 and h-2 denote divergence times (in coalescent units), *ti* parameters denote intermixture times, *mti* parameters denote start/end of migration epochs, γ is the inheritance probability, and mr is the population migration rate3 (see main text).

For each simulation setting, we simulated 20 data sets with 200 1-kb loci (in this part, we did not vary the sequence lengths and numbers of loci). We set the population mutation rate at 0.02 across all the branches. Furthermore we set the inheritance probability *γ* and the migration rate mr each to 0.20 (here, *mr = 2Nm*, where N is the effective population size, and m is the fraction of the recipient population that is made up of migrants from the donor population in each generation). We set *h*_1_ = 9, *h*_2_ = 6. For the intermixture model (Fig. 17(a)), we set *t*_2_ = 3, and varied (*t*_1_,*t*_3_) to take on the values (4,2), (5,1), and (6,0) so that the elapsed time, denoted by Δt, between subsequent reticulation events is 1, 2, or 3. For the gene flow model (Fig. 17(b)), we set (*mt*_1_,…,*mt*_6_) to (6,5,3.5,2.5,1,0), so that the duration of each gene flow epoch is 1 and the time elapsed between between two consecutive epochs, denoted by Δ*mt*, is 1.5. The commands for the ms and Seq-gen programs are given in Supplementary Materials.

For each data set, we ran an MCMC chain of 8 × 10^6^ iterations with 1 × 10^6^ burn-in. One sample was collected from every 5,000 iterations, resulting in a total of 1,400 collected samples. We summarized the results based on 28,000 samples from 20 replicates for each parameter setting.

Table 1 shows the population mutation rates, divergence times, and numbers of reticulations estimated by our method on data generated under the models of Fig. 17(a) and Fig. 17(b). As the results show, the method performs very well in terms of estimating the divergence times and population mutation rates, regardless of whether the data were generated under an intermixture model or a gene flow model. Furthermore, for these two parameters, the estimates are stable while varying the elapsed times between consecutive reticulation events.

**TABLE 1.** Estimated population mutation rates (θ), divergence times (h_1_ and h_2_), and numbers of reticulations (#reti) as a function of varying Δ*t* in the model of Fig. 17(a) and Δ*mt* in the model of Fig. 17(b). The divergence times were estimated in units of expected number of mutations per site and are reported in coalescent units by dividing by *θ*/2 = 0.01.

| Case | $\theta$ | $h_1$ | $h_2$ | $\#reti$ |
| --- | --- | --- | --- | --- |
| $\Delta t = 1$ | $2.2 \pm 0.2e^{-2}$ | $8.9 \pm 0.1$ | $5.9 \pm 0.1$ | $1.2 \pm 0.4$ |
| $\Delta t = 2$ | $2.2 \pm 0.2e^{-2}$ | $8.9 \pm 0.1$ | $5.9 \pm 0.1$ | $2.0 \pm 0.0$ |
| $\Delta t = 3$ | $2.1 \pm 0.3e^{-2}$ | $9.0 \pm 0.1$ | $6.0 \pm 0.1$ | $2.6 \pm 0.5$ |
| $\Delta mt = 1.5$ | $2.3 \pm 0.3e^{-2}$ | $8.9 \pm 0.1$ | $6.0 \pm 0.1$ | $2.1 \pm 0.3$ |

As for the estimated number of reticulations, it becomes more accurate as the elapsed times between consecutive reticulations is larger. To better understand the factors that affect the detectability of reticulations, we plotted histograms of the true and estimated coalescence times of the most recent common ancestor (MRCA) of alleles from *B* and *C* in Fig. 18. Here, the true coalescence times are obtained from the true gene tree simulated generated by the program ms. The estimated coalescence times are sampled by our method along with the gene tree topologies. For the estimated coalescence times, we plot them based on all the collected samples, which is why the histograms of estimated coalescence times are smoother than those of the true ones.

**FIGURE 18.**
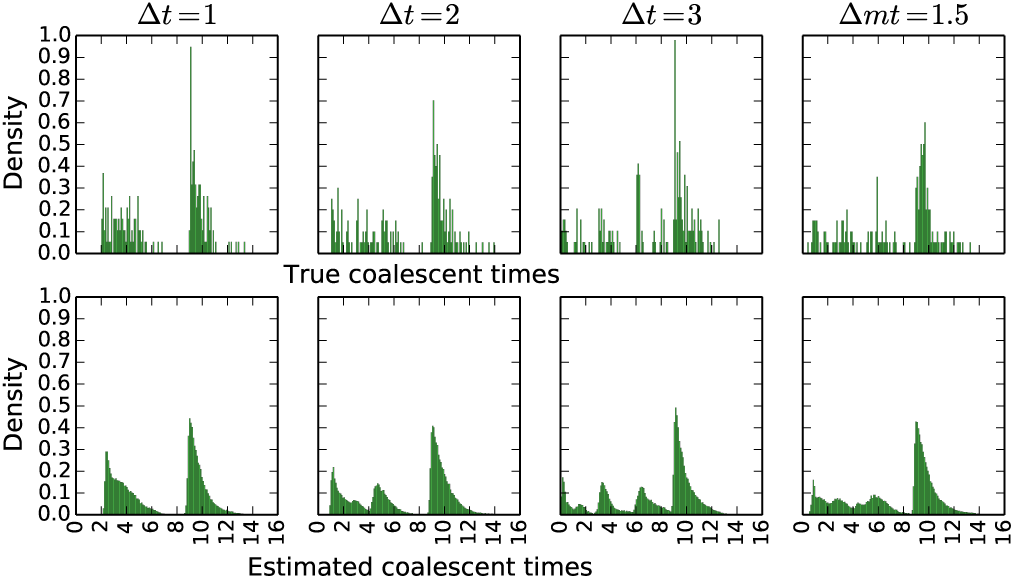
Histograms of the true (top) and estimated (bottom) coalescence times (in coalescent units) of the MRCA of alleles from *B* and *C* on data generated under the models of Fig. 17(a) and Fig. 17(b).

As Fig. 17(a) and Fig. 17(b) show, the coalescence times of alleles from B and C would form a mixture of four distributions: three due to the three reticulation events, and one above the root of the phylogenetic network. As the left three columns of panels in Fig 18 show, under an intermixture model, as δ*t* increases, the signal for a mixture of four distributions of (A,B) coalescence times becomes much stronger, thus pointing to three reticulations in addition to the coalescence events above the root of the phylogeny. This is why, under the intermixture model, the method’s performance in terms of the estimated number of reticulations improves as Δt increases. However, on data simulated the under the gene flow model (the rightmost column of panels in Fig. 18), the signal of the mixture of four distributions of *(A,B)* coalescence times is surprisingly stronger than that under the intermixture model with the comparable Δt =1 and Δt = 2.

Fig. 19 shows results similar to those reported in Fig. 18, with the only difference being that these are the coalescence times from all 4,000 loci generated from the 20 data sets of 200 loci each. Effectively, this is the signal in a data set of 4,000 independent loci. Clearly,9 the signal is much stronger than in data sets of 200o loci, and all reticulations would be recoverable under the intermixture model for Δt = 2,3 and for the gene flow model.

**FIGURE 19.**
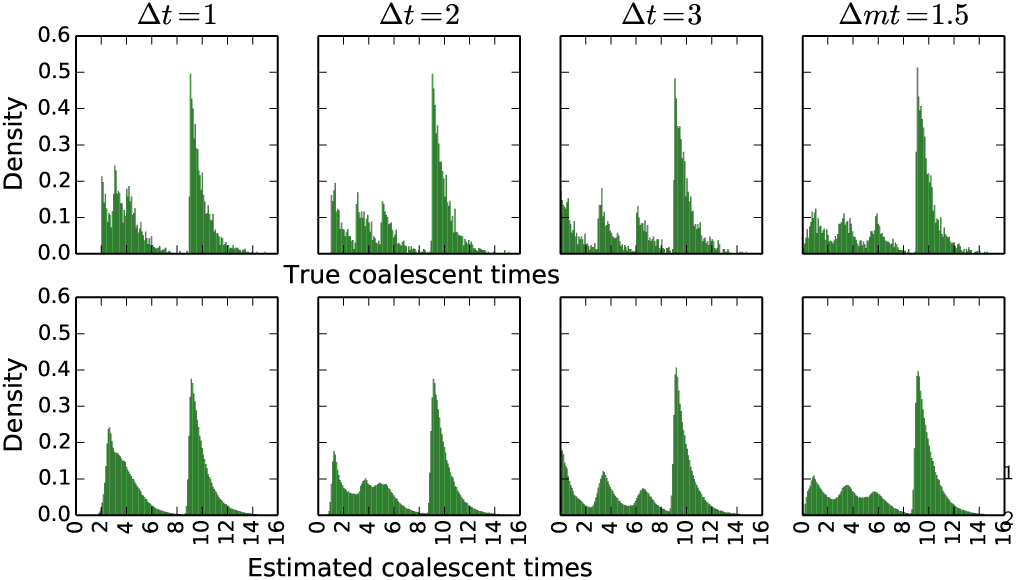
Histograms of the true (top) and estimated5 (bottom) coalescence times (in coalescent units) of the MRCA of alleles from *B* and *C* on 4,000 loci generated under the models of Fig. 17(a) and Fig. 17(b).

We also ran simulations where we varied the number of individuals sampled from species B (we sampled 1, 3, and 5 individuals). The results improve as the number of individuals increases from 1 to 3, but no discernible improvement is achieved under our simulation settings when the number of individual is increased to 5. Results are given in the Supplementary Materials.

To assess the performance of our method on the simpler case of a single reticulation event, we considered the networks in Fig. 17(c) and Fig. 17(d), set *h*_*1*_ =2.5, *h*_*2*_ = 1.5, and *mt*_*1*_ = *h*_*2*_, and varied *t,mt*_*2*_ ∈{1,0}. As the results in Table 2 demonstrate, our method estimated the population mutation rate θ, the divergence times *h*_*1*_ and *h*_*2*_, and the inheritance probability/migration rate very accurately under all cases. The method did very well also in terms of estimating *t* and *mt*_*2*_; results in Supplementary Materials.

**TABLE 2.** Estimated population mutation rates (0), divergence times (*h*_1_ and h_2_), inheritance/migration rates, and numbers of reticulations (#reti) as a function of varying *t* in the model of Fig. 17(c) and *mt*_*2*_ in the model of Fig. 17(d). The divergence times were estimated in units of expected number of mutations per site and are reported in coalescent units by dividing by θ/2 = 0.01.

| Case | $\theta$ | $h_1$ | $h_2$ | $\gamma$ ($mr$) | $\#reti$ |
| --- | --- | --- | --- | --- | --- |
| $t=1$ | $2.0 \pm 0.2e^{-2}$ | $2.5 \pm 0.1$ | $1.5 \pm 0.1$ | $0.20 \pm 0.05$ | $1.0 \pm 0.0$ |
| $t=0$ | $2.0 \pm 0.2e^{-2}$ | $2.5 \pm 0.1$ | $1.5 \pm 0.1$ | $0.21 \pm 0.04$ | $1.0 \pm 0.0$ |
| $mt_2=1$ | $2.0 \pm 0.2e^{-2}$ | $2.5 \pm 0.1$ | $1.5 \pm 0.1$ | $0.18 \pm 0.05$ | $1.0 \pm 0.0$ |
| $mt_2=0$ | $2.2 \pm 0.2e^{-2}$ | $2.5 \pm 0.1$ | $1.5 \pm 0.1$ | $0.17 \pm 0.04$ | $1.0 \pm 0.0$ |

A single reticulation was detected for all cases of 3 intermixture and gene flow. We plotted the histograms of 4 the true and estimated coalescence times of the MRCA of alleles from *B* and *C* in Fig. 20. As the figure shows, the distributions of estimated coalescence times match the distributions of true coalescence times very well. Furthermore, when using 4, 000 loci, the signal becomes even stronger; results in Supplementary Materials.

Finally, we assessed the performance of our method on cases where the reticulation event involves sister taxa. Fig. 17(e) and Fig. 17(f) show the cases we considered, with setting *h*_*1*_ = 2.5 and *h*_2_ = 1.5, and varying *t,mt∈* {1,0}.

As the results in Table 3 demonstrate, our method obtained very accurate estimates of the various parameters under *t* = 0 and *mt* = 0. Under the cases of intermixture with *t* = 1 and gene flow with *mt* = 1, our method did not detect the reticulation, which resulted in an underestimation of *h*_2_. In the case of *mt* = 0, the migration rate was severely underestimated, most likely due to the short time interval between the migration and divergence events between *A* and *B*. The method did very well also in terms of estimating *t* and *mt*; results in Supplementary Materials.

**FIGURE 20.**
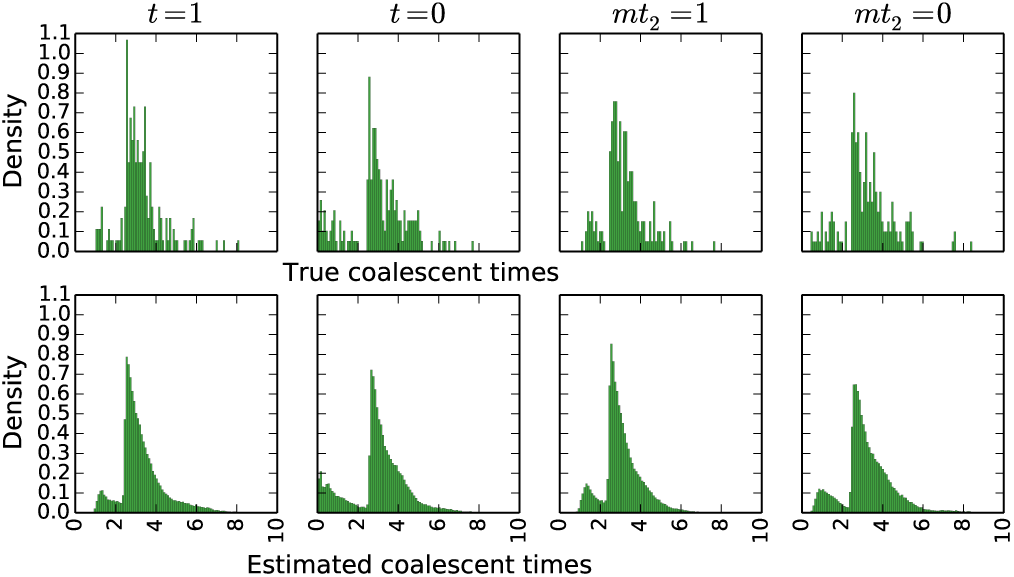
Histograms of the true (top) and estimated (bottom) coalescence times (in coalescent units) of the MRCA of alleles from *B* and *C* on data generated under the models of Fig. 17(c) and Fig. 17(d).

**TABLE 3.** Estimated population mutation rates (*θ*), divergence times (*h*_1_ and *h*_2_), inheritance/migration rates, and numbers of reticulations (#reti) as a function of varying *t* in the model of Fig. 17(e) and *mt* in the model of Fig. 17(f). The divergence times were estimated in units of expected number of mutations per site and are reported in coalescent units by dividing by *θ /*2=0.01.

| Case | $\theta$ | $h_1$ | $h_2$ | $\gamma$ | #reti |
| --- | --- | --- | --- | --- | --- |
| $t=1$ | $2.0 \pm 0.2e^{-2}$ | $2.5 \pm 0.1$ | $1.3 \pm 0.1$ | NA | $0.0 \pm 0.0$ |
| $t=0$ | $2.0 \pm 0.2e^{-2}$ | $2.5 \pm 0.1$ | $1.5 \pm 0.0$ | $0.21 \pm 0.06$ | $1.0 \pm 0.0$ |
| $mt=1$ | $2.0 \pm 0.2e^{-2}$ | $2.5 \pm 0.1$ | $1.4 \pm 0.1$ | NA | $0.0 \pm 0.0$ |
| $mt=0$ | $2.2 \pm 0.2e^{-2}$ | $2.5 \pm 0.1$ | $1.5 \pm 0.1$ | $0.11 \pm 0.06$ | $1.0 \pm 0.0$ |

We plotted the histograms of the true and estimated coalescence times of the MRCA of alleles from *A* and *B* in Fig. 21. When *t* = 1 and *mt* = 1, the signal of reticulation is very low, which explains the failure of our method to detect it. In the cases of *t* = 0 and *mt* = 0, the distributions of estimated coalescence times match those of true coalescence times very well. When using 4,000 loci, the signal becomes even stronger; results in Supplementary Materials.

**FIGURE 21.**
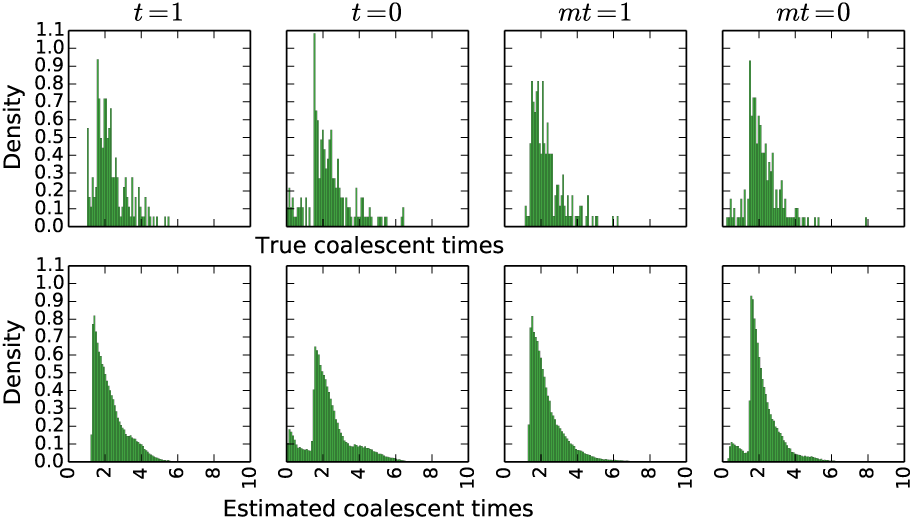
Histograms of the true (top) and estimated (bottom) coalescence times (in coalescent units) of the MRCA of alleles from *A* and *B* on data generated under the models of Fig. 17(e) and Fig. 17(f).

### Analysis of a 106-locus Yeast Data Set

The yeast data set of (Rokas *et al.,* 2003) consists of 106 loci from seven Saccharomyces species, *S. cerevisiae* (Scer), *S. paradoxus* (Spar), *S. mikatae* (Smik), *S. kudriavzevii* (Skud), *S. bayanus* (Sbay), *S. castellii* (Scas), *S. kluyveri* (Sklu). Rokas *et al.* (Rokas *et al.,* 2003) reported on extensive incongruence of single-gene phylogenies and revealed the species tree from concatenation method (Fig. 22(a)). Edwards *et al.* (Edwards *et al.,* 2007) reported as the two main species trees and gene tree topologies sampled from BEST (Liu, 2008) the two trees shown in Fig. 22(a-b). The other gene tree topologies (Fig. 22(c)) exhibited weak phylogenetic signals among Sklu, Scas and the other species. Bloomquist and Suchard (Bloomquist and Suchard, 2010) reanalyzed the data set without Sklu since it added too much noise to their analysis. Their analysis resulted in many horizontal events between Scas and the rest of the species because the Scas lineage-specific rate variation is much stronger than that of the other species. Yu *et al.* (Yu *et al.,* 2013b) analyzed the 106-locus data set restricted to the five species Scer, Spar, Smik, Skud, and Sbay and identified a maximums parsimony network that supports a hybridization from Skud to Sbay with inheritance probability of 0.38.

**FIGURE 22.**
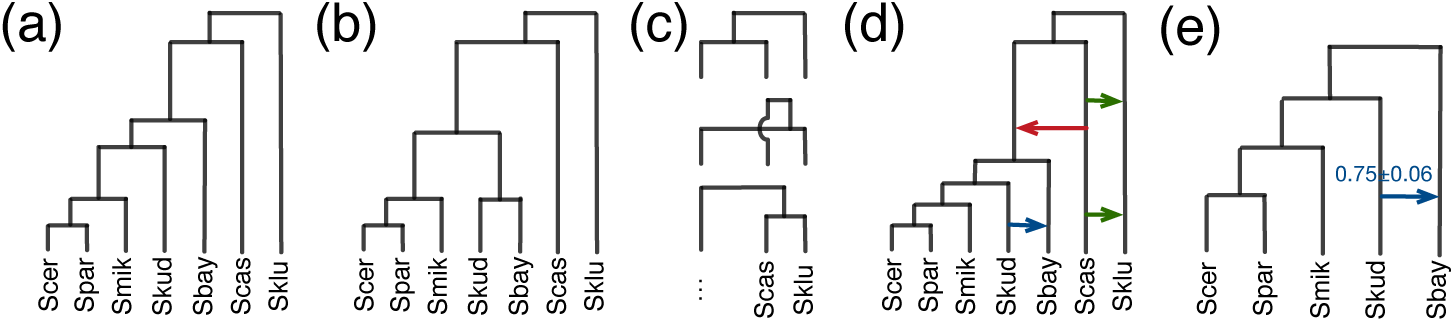
Results on the yeast data set of (Rokas *et al.,* 2003). (a) The species tree inferred using the concatenation method (Rokas *et al.,* 2003) and the main species tree and gene tree topology sampled using BEST (Edwards *et al.,* 2007). (b) The second most frequently sampled species and gene tree topology by BEST (Edwards *et al.,* 2007). (c) Many other gene tree topologies were sampled by BEST (Edwards *et al.,* 2007), indicating weak phylogenetic signals among Sklu, Scas, and the rest of the species. (d) The MAP phylogenetic network inferred by our method on all 106 loci. (e) The single phylogenetic network inferred using all 106 loci from the five species Scer, Spar, Smik, Skud, Sbay.

Analyzing the 106-locus data set using our method, the 95% credible set contains many topologies with similar hybridization patterns; the representative network is shown in Fig. 22(d). All the previous findings are encompassed by the networks inferred by our method. The two hybridizations between Sklu and Scas (green edges in 22(d)) indicate the weak phylogenetic signals among Sklu, Scas and the rest of the species. The hybridization from Scas to the other species except for Sklu (red edge in 22(d)) captures the stronger lineage-specific rate variation in Scas. Finally, the hybridization from Skud to Sbay (blue edge in 22(d)) resolves the incongruence between the two main species tree topologies in 22(a-b).

We then analyzed the 106-locus data set restricted to the five species Scer, Spar, Smik, Skud, and Sbay. The phylogenetic signal in this data set is very strong—the consensus trees of 99 out of the 106 loci contain two internal branches. The MPP phylogenetic network in Fig. 22(f) contains the hybridization from Skud to Sbay, which is identical to the sub-network in Fig. 22(d). See Supplementary Materials for full details. In summary, analysis of the yeast data set demonstrates the effect of phylogenetic signal in the individual loci on the inference and the care that must be taken when selecting loci of analysis of reticulate evolutionary histories.

We compared these analyses to ones obtained by the method of (Wen *et al.* 2016a) when the input data consist of gene tree estimates. When the gene tree estimates on all seven Saccharomyces species are used, the 95% credible set consisted of a single network that is shown in Fig. 22(d), yet with only the single reticulation from Skud to Sbay. When the gene tree estimates on the subset of five species were used as input, the 95% credible set consisted of a single network that is shown in Fig. 22(e), in agreement with the results based on co-estimation from the sequence data directly.

Finally, we quantified the Robinson-Foulds distances between the locus-specific gene tree estimates obtained by our method and by RAxML. The distances were 0.33±0.19 for the 7-taxon data set, and 0.33±0.16 for the 5-taxon data set. It is worth noting that these distances are very similar to those observed in Fig. 12 above. Full details and further results for this data set are given in Supplementary Materials.

## DISCUSSION

To conclude, we have devised a Bayesian framework for sampling the parameters of the MSNC model, including the species phylogeny, gene trees, divergence times, and population sizes, from sequences of multiple independent loci. Our work provides the first general framework for Bayesian phylogenomic inference from sequence data in the presence of hybridization. The method is publicly available in the open-source software package PhyloNet (Than *et al.* 2008). We demonstrate the utility of our method on simulated data and three biological data sets.

Our results demonstrate several important aspects. First, ignoring hybridization when it had occurred results in underestimating the divergence times of species and overestimating the coalescence times of individual loci. Second, co-estimation of species phylogeny and gene trees results in more accurate gene tree estimates than the inferences of gene trees from sequences directly. Third, comparing to existing phylogenetic network inference methods (Wen *et al.* 2016a; Yu *et al.* 2014) that use gene tree estimates as input, our method not only estimates more parameters, such as divergence times and population sizes, but also estimates more accurate phylogenetic networks from fewer loci. Further, we assessed the performance of our model and method on simulated data generated under a gene flow model. Our method performed very well on such data. However, given the nature of our abstract phylogenetic network model, a gene flow epoch is estimated as a single reticulation event. Finally, we analyzed a 106-locus yeast data set and demonstrated for empirical data the differences in results one obtains when co-estimating the gene and species phylogenies when compared to inferences from gene tree estimates.

Finally, we identify several directions for further improvements of our proposed approach. First, while priors on species trees, such as the birth-death model, have been developed and employed by inference methods, similar prior distributions on phylogenetic networks are currently lacking. Second, while techniques such as the majority-rule consensus exist for summarizing the trees sampled from the posterior distribution, principled methods for summarizing sampled networks are needed. Last but not least, the sequence data used here, and in almost all phylogenomic analyses, consist of haploid sequences of randomly phased diploid genomes. The effect of random phasing on inferences in general needs to be studied in detail. Furthermore, the model could be extended to work directly on unphased data by integrating over possible phasings (Gronau *et al.* 2011).

## SUPPLEMENTARY MATERIAL

Supplementary material, including data files and online-only appendices, can be found in the Dryad data repository at.

